# Coronary artery disease is linked with demyelination and iron deposition in white matter watershed areas

**DOI:** 10.64898/2026.03.03.709359

**Authors:** Ali Rezaei, Zacharie Potvin-Jutras, Stefanie A. Tremblay, Safa Sanami, Dalia Sabra, Julia Huck, Christine Gagnon, Lindsay Wright, Ilana R. Leppert, Christine L. Tardif, Josep Iglesies-Grau, Anil Nigam, Louis Bherer, Claudine J. Gauthier

**Author notes:** **Correspondence to**: Claudine J. Gauthier, **Full address**: 7141, rue Sherbrooke Ouest, Montreal, Quebec H4B 1R6, Canada, **E-mail:**.

## Abstract

Coronary artery disease increases risk of cognitive decline and stroke and is associated with white matter alterations. However, the biological basis of these changes remains unclear. Myelin content and iron deposition are crucial measures of white matter health and can be measured with quantitative MRI. This study investigated whether myelin and iron alterations occur in coronary artery disease, and their relationship with cognition.

In this cross-sectional study, 46 individuals with coronary artery disease and 40 healthy controls aged > 50 years, with normal cognition underwent 3T MRI and cognitive assessments. Quantitative MRI metrics (susceptibility, magnetization transfer saturation, *R*_2_^∗^ and *R*_1_ relaxation rates) were calculated in the border zones between adjacent arterial territories (watershed regions) and in the areas outside these borders (non-watershed regions).

Relative to controls, the coronary artery disease group showed lower myelin and higher iron content, as measured by lower magnetization transfer saturation and *R*_1_, and higher susceptibility specifically in watershed regions. Importantly, these microstructural alterations were associated with poorer cognitive performance in the coronary artery disease group with lower magnetization transfer and *R*_1_related to poorer global cognition and with higher magnetic susceptibility with poorer verbal memory.

These findings suggest that coronary artery disease is associated with demyelination and iron deposition in white matter, most prominently in watershed regions, which are known for their susceptibility to stroke. The association of these microstructural alterations with cognition highlights the role of white matter as a key vulnerable region and a promising focus for future mechanistic and therapeutic studies.

## Introduction

Heart diseases are the leading causes of morbidity and mortality worldwide, taking 17.9 million lives each year [1]. Coronary artery disease (CAD), which is caused by the buildup of atherosclerotic plaque within the coronary arteries that feed the heart, is the most prevalent form of heart disease [2]. CAD, while primarily a cardiac condition, is increasingly recognized to have detrimental effects beyond the heart. Recent studies show that individuals with CAD are at greater risk of cognitive decline [3, 4] and dementia [5]. In addition, individuals with CAD show poorer cognitive performance [6]. The underpinnings of these cognitive effects are likely to be complex and multifactorial, but changes in white matter (WM) are emerging as important contributors. WM health plays a crucial role in maintaining cognitive function, both in healthy aging [7] and pathological conditions such as small vessel disease [8] and CAD [9]. WM health is often assessed by the burden of white matter hyperintensities (WMH), a measure of vascular lesion load, and WMH burden has been consistently associated with poorer cognitive performance [10–12]. However, emerging evidence suggests that WMH captures only the tip of the iceberg of WM degradation and that the normal appearing white matter (NAWM) also undergoes more subtle microstructural alterations that contribute to cognitive dysfunction [13–15]. Indeed, studies in stroke [16, 17] and cerebral small vessel disease [18] have demonstrated that NAWM integrity plays a critical role in preserving cognitive function.

The brain is supplied by several arteries, including the anterior (ACA), middle (MCA), posterior cerebral (PCA) and vertebrobasilar (VB) arteries, each with its own territory. At the border zones of these territories lie the watershed regions. Because these watershed regions lie at the limit of each main arterial territory and are perfused by smaller terminal arteries, they more frequently experience low perfusion pressure [19] and are especially vulnerable to stroke [20]. Thus, watershed regions are vulnerable to hypoperfusion and therefore transient ischemia, leading to more frequent development of WMH than in other brain regions [21]. Given that CAD is linked to vascular dysfunction and hypoperfusion [22], it is plausible that CAD also affects NAWM microstructure, particularly in vulnerable watershed regions. However, this has not been investigated.

WM microstructure is complex, but determined in large part by myelin and iron which can be assessed using quantitative MRI (qMRI) techniques [23]. Reduced myelin content has been observed in severe heart diseases including heart failure and congenital heart disease [24, 25] and is linked to poorer global cognition [26, 27]. Furthermore, studies have shown that vascular diseases including stroke are also associated with increased neuroinflammation which in turn has been linked to increased activity of iron-rich microglia activity [28] and to cognitive decline [29]. In CAD, while some studies showed WM microstructural alterations, they have largely focused on metrics of axonal structure [30–32], with a recent added focus on myelin [9]. There are currently no studies on whether iron content is altered in CAD, and whether iron and myelin changes co-occur. Therefore, we do not currently know the contribution of myelin loss and inflammatory processes on the effects of CAD on WM microstructural health. Here, we will assess both, as well as important considerations related to the spatial distribution of these microstructural deficits in the watershed regions of the brain.

In this cross-sectional study, we explore NAWM microstructure using qMRI within watershed and non-watershed regions, in individuals with CAD compared to healthy controls. We hypothesize that individuals with CAD will have lower myelin content and higher iron deposition than healthy controls, and that these effects will be exacerbated in watershed areas. Finally, we expect lower myelin and higher iron to be linked to poorer cognitive performance in individuals with CAD. Understanding these relationships will offer novel insight into the underlying mechanisms by which vascular pathology contributes to cognitive decline and may ultimately provide therapeutic targets for interventions in individuals with CAD.

## Materials and methods

### Participants

Ninety nine (99) participants of 50 years and above were recruited, from which 86 completed the study (46 individuals with CAD; CAD group, and 40 healthy controls; HC group). Out of 9 participants that discontinued, five participants did not complete the study due to interruptions during the COVID-19 pandemic, one participant was excluded due to claustrophobia, and three participants chose not to participate for personal reasons (e.g., loss of interest, issues with scheduling). Because the MRI acquisition also included a hypercapnia manipulation (not used here [22], which can cause discomfort in some individuals, our attrition rate was higher than that of conventional MRI sessions. Participants who had all the required MRI data were included in this study (N= 90). However, three participants were ultimately excluded due to the presence of severe motion artifacts in MRI data (N=3) and one was found to have an incidental finding in their MRI data (N=1), resulting in a sample size of 86. Of those, 46 had CAD (age = 68.2 ± 9 years, 8 females) and 40 were HC (age = 65.5 ± 8, 11 females). Fig. 1 shows the flowchart of participants’ inclusion.

**Figure 1.**
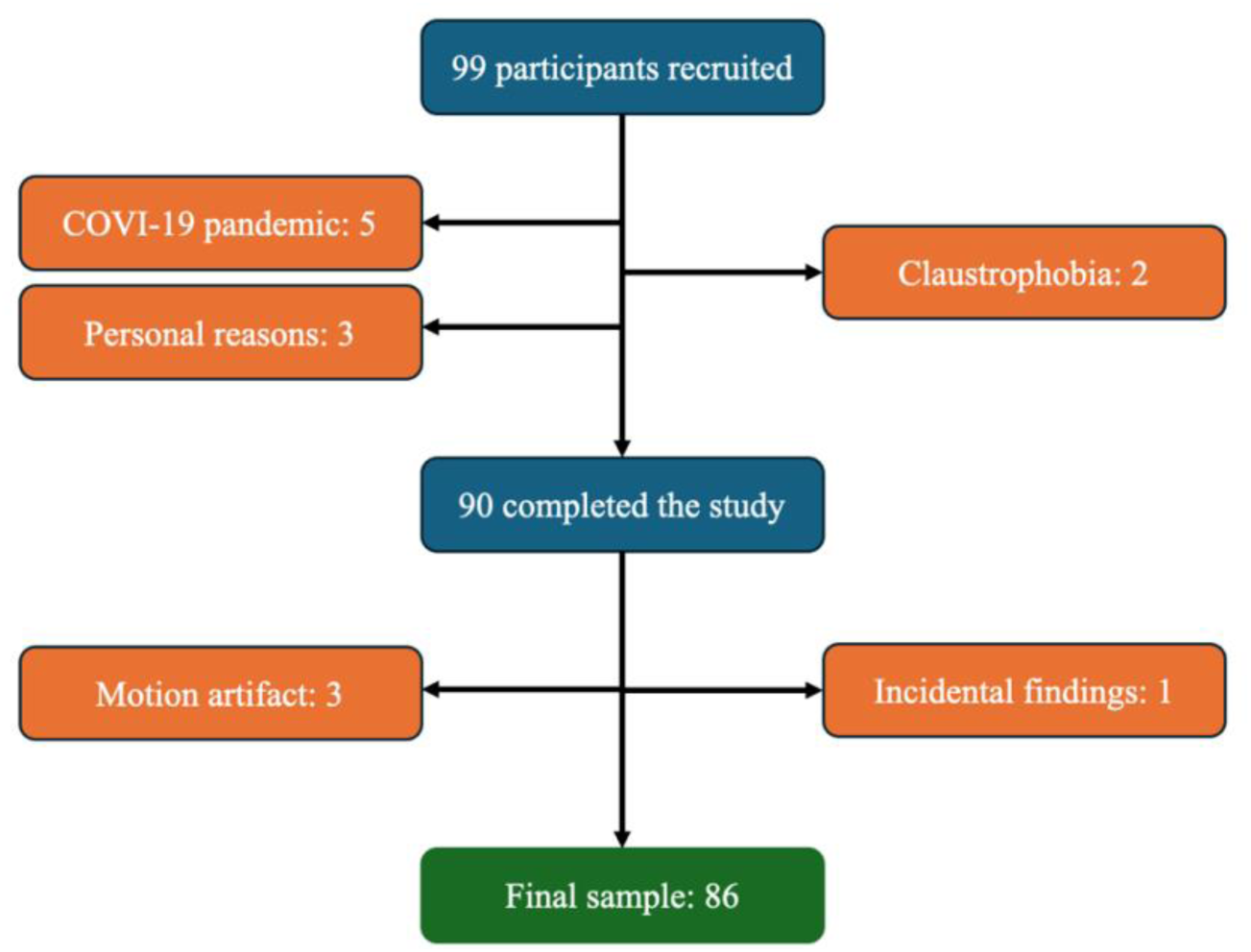
Flowchart of participants inclusion. A total of 99 participants were screened and consented. Of these, 9 discontinued their participation, leaving 90 who completed the study. Among them, 3 were excluded due to motion artifacts in gradient echo data and 1 was excluded due to incidental findings, resulting in a final analyzed sample of 86 participants.

The study was approved by the Comité d’éthique de la recherche et du développement des nouvelles technologies (CÉRDNT) de l’Institut de Cardiologie de Montréal, in accordance with the Declaration of Helsinki. Written informed consent was obtained at the first visit. A health questionnaire, which had been previously filled out over the phone to determine eligibility, was also reviewed at this visit. Finally, the mini-mental state examination (MMSE) was administered to ensure eligibility. All participants then completed a neuropsychological battery and an MRI acquisition during separate sessions.

Inclusion criteria for our CAD group included documented coronary artery disease (prior acute coronary syndrome, prior coronary angiography or revascularization, or myocardial ischemia documented on myocardial scintigraphy). All participants had to be fluent in either English or French (for informed consent and cognitive assessment).

Exclusion criteria for all participants included history of stroke, neurological, psychiatric or respiratory disorders, thyroid disease, potential cognitive impairment (MMSE < 25), tobacco use, high alcohol consumption (more than 2 drinks per day), contraindications for MRI (e.g., ferromagnetic implant, claustrophobia), and use of oral or patch hormone therapy. Participants were also excluded if they had surgery under general anesthesia within the last 6 months, a recent acute coronary event (< 3 months), chronic systolic heart failure (resting left ventricular ejection fraction < 40%), symptomatic aortic stenosis, severe nonrevascularizable coronary artery disease, including left main coronary stenosis, awaiting coronary artery bypass surgery, implanted automatic defibrillator or permanent pacemaker. Excessive discomfort due to hypercapnia (> 5 on the dyspnea scale of Banzett [33] also constituted an exclusion criterion [22] though this data is not used here. Participants in the HC group had to be free of any cardiac and neurological issues, diabetes, hypertension, and current use of medications known to be vasoactive (e.g., beta-blockers).

### MRI protocols

MRI data were acquired on a 3T Siemens Magnetom Skyra scanner at the Montreal Heart Institute with a 32-channel head coil. High-resolution T_1_-weighted structural images with a Magnetization Prepared RApid Gradient Echo (MPRAGE) sequence (TR= 2300 ms, TE= 2.32 ms, flip angle= 8°, resolution= 0.9 mm isotropic) and T_2_-weighted axial fluid attenuated inversion recovery (FLAIR) images (TR= 9000 ms, TE= 91 ms, TI= 2500 ms, flip angle= 150°, resolution = 0.9 x 0.9 x 5.0 mm) were acquired for brain tissue and lesion segmentation. A 3D multi-echo gradient echo (ME-GRE) (TR/TE1/TE2/TE3/TE4/flip angle = 20 ms/6.92 ms/13.45 ms/19.28 ms/26.51 ms/9°, resolution = 0.7 × 0.7 × 1.4 mm) phase and magnitude data were acquired for all the coil channels separately for the reconstruction of quantitative susceptibility maps. Flow compensation of the first time of echo (TE) data was done by nulling the gradient moment to minimize flow artifact effects for the venous mapping [34].

Three gradient echo sequences (TR = 33 ms, TE = 3.81 ms, flip angle = 10°, resolution = 2 mm isotropic), one with (MT-w) and one without a preparatory MT pulse (MT-off), and a T1w image (TR = 15 ms, TE = 3.81 ms, flip angle = 25°, resolution = 2 mm isotropic) were acquired for MTsat computation. An off-resonance MT pulse (off-resonance frequency = 2.2 kHz, duration = 12.8 ms, flip angle = 540°) was applied prior to RF excitation to obtain MT-weighting [35]. A B1 map (an anatomical image and a flip angle map) (TR = 19780 ms, TE = 2.36 ms, flip angle = 8°, resolution = 3.1 × 3.1 × 8 mm) was also acquired using a fast gradient echo sequence (TurboFLASH) with a preconditioning radiofrequency pulse to correct B1 field inhomogeneities [36].

### MRI data analysis

#### Tissue segmentation

Preprocessing of the T_1_-weighted images included brain extraction via *brain extraction based on nonlocal Segmentation Technique* (BEaST) [37], denoising, intensity nonuniformity correction, and intensity normalization correction, all performed using the MINC toolkit [38]. Tissue compartments, including cerebrospinal fluid, grey matter, white matter and white matter hyperintensities were segmented from the T_1_-weighted and FLAIR images using the automated Brain tIssue SegmentatiON (BISON) random forest classifier [39].

#### Quantitative Susceptibility Mapping (QSM) reconstruction

To reconstruct the QSM maps, ME-GRE magnitude data from the individual channels were combined using the square root of the sum of squares for each TE [40]. The multi-channel ME-GRE phase data was combined and unwrapped using the ROMEO toolbox to ensure spatiotemporal coherence and temporal stability [41]. The channel-combined and unwrapped phase data of all TEs were combined into a single phase map using the MEDI toolbox non-linear complex fitting method [42]. Then, using the first echo and echo-combined phase files, two QSM (*χ*) maps were reconstructed using Total Generalized Variation (TGV) QSM reconstruction [40]. This method was selected to minimize noise propagation through consecutive QSM processing steps and preserving edges between different tissue types, especially the edges between veins and surrounding tissue. To remove the effect of paramagnetic deoxyhemoglobin in veins on susceptibility measurements in regional tissues, the veins were extracted using a multi-scale recursive ridge filtering method [43] from the QSM data reconstructed from the first TE data, since this was the only echo time with flow compensation. The rest of the assessments was done using the QSM map reconstructed from the echo-combined phase. All the QSM maps were zero-referenced by the mean *χ* values of the cerebrospinal fluid in the ventricles [44].

### R_2_^∗^ relaxation rate map reconstruction

To reconstruct *R*_2_^∗^ from the ME-GRE magnitude data, we first corrected the signal for macroscopic field inhomogeneity [45]. The B0 field map was calculated by applying linear fitting to the unwrapped phase data generated during QSM processing. The background field was estimated from the total field using regularization-enabled sophisticated harmonic artifact reduction for phase data with varying kernel sizes implemented in the MEDI toolbox [42, 45, 46]. Then, The *R*_2_^∗^ value was fitted from the corrected magnitude data to the mono-exponential *R*_2_^∗^ decay using auto-regression on linear operations (ALRO) using the MEDI toolbox [47].

### MTsat and R_1_ relaxation rate map reconstruction

The MTsat and *R*_1_maps were calculated using the hMRI-toolbox (v0.3.0) [48]. The *B*_1_ transmit bias-corrected field was computed using an anatomical map and a scaled flip angle map using the “Create hMRI maps” module. Then, the *B*_1_ correction field, MT-w, MT-off and T_1_-weighted images were included in the “Create hMRI maps” module to estimate *R*_1_ maps with default parameters. Finally, non-brain voxels were removed from the *R*_1_ maps using FSL’s brain extraction tool [49].

#### Registration

Registration of the vascular territories atlas from the standard Montreal Neurological Institute (MNI) space and segmentation files to the subjects’ native QSM space were done using *advanced normalization tools* (ANTs) (https://github.com/ANTsX/ANTs). For the registration of MPRAGE data to MNI space, a multistage approach that combined rigid, affine, and nonlinear (SyN) transformations with a multi-resolution strategy was used. For the co-registration of subjects’ MPRAGE data and *R*_1_ map to native QSM space, a combined rigid and affine transformation was utilized.

#### Definition of Regions of Interest (ROIs)

White matter hyperintensities were excluded from the analysis by excluding the lesion mask generated using the BISON toolbox from the WM mask to generate a NAWM mask for each participant. In addition, veins were removed from the NAWM mask by excluding the vein mask derived from the multiscale vessel filter. The resultant NAWM mask was eroded by two voxels to avoid the regions close to the border of the ventricles and the grey matter as well as white matter hyperintensities.

Individualized vascular ROIs were created by combining a cerebral arterial territory atlas [50] with individualized NAWM masks (which excluded veins and WMHs). In addition, the watershed regions (WS) were defined by dilating each arterial territory by 3.5 mm (5 voxels) and using the intersection of the interface between adjacent territories. Therefore, the intersection of the dilated ACA and MCA NAWM regions is the ACA_MCA watershed territory, the intersection of dilated MCA and PCA is the MCA_PCA watershed, and PCA and VB is the PCA_VB watershed (Figure 2). The total watershed region includes the combination of all individual WS regions. The voxels within the NAWM of each arterial territory that are not within these watershed regions constitute the non-watershed regions (NWS). These ROIs were used to extract regional averages for each parameter in the WS and NWS regions. Fig. 2 shows an example of NAWM in WS and NWS regions overlaid on the susceptibility (*χ*), *R*_2_^∗^, *R*_1_ and MTsat data of one of the participants.

**Figure 2.**
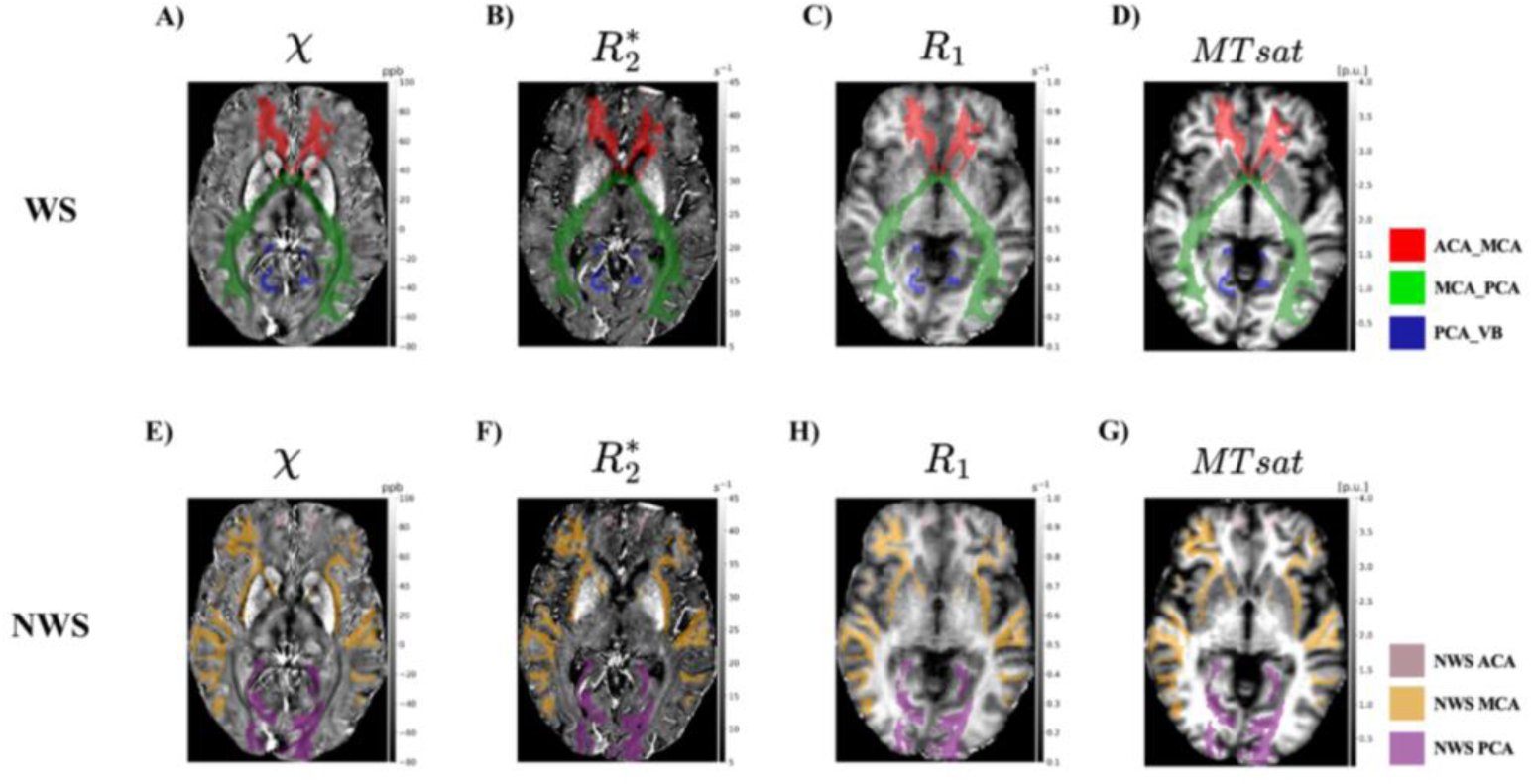
Individualized WS and NWS regions overlaid on example qMRI images in QSM native space. **A** and **E**: χ, **B** and **F**: R_2_^∗^, **C** and **H**: R_1_ and **D** and **G**: MTsat. The top row shows the WS regions in which the red color represents the WS region between ACA and MCA arterial territories (ACA_MCA), the green color represents the WS region between MCA and PCA arterial territories (MCA_PCA) and the blue color represents the WS region between PCA and VB arterial territories (PCA_VB).The bottom row shows the NWS regions in which the pink color represents the NWS region of ACA arterial territory (NWS ACA), the orange color represents the NWS region of MCA arterial territory (NWS MCA) and the purple color represents the NWS region of PCA arterial territory (NWS PCA). The unit for χ is part per billion (ppb). The unit for R_2_^∗^ and R_1_ is s^-1^. The unit for MTsat is percent units (p.u.).

#### Cognitive Assessment

A comprehensive neuropsychological assessment was administered in the following order: Global cognition was evaluated using the Montreal Cognitive Assessment (MoCA) [51]. The Rey Auditory Verbal Learning Test (RAVLT) was used to assess verbal episodic memory, where participants recalled words from a 15-word list across 5 learning trials, immediately after an interference list, and again after a 30-minute delay. The Delis-Kaplan Executive Function System (D-KEFS) Color Word Interference Test (CWIT) was used to assess executive function, which includes four conditions: color naming, reading, inhibition, and switching [52].

All scores were transformed into standardized z-scores and 2 composite scores were then created using those z-scores: (1) executive functioning = (TMT B+ CWIT 3+ CWIT 4 scores)/3); and (2) verbal episodic memory (immediate recall + delayed recall + total words scores recalled during the 5 learning trials from the RAVLT test)/3).[53] Z-scores of timed tests were multiplied by −1 so that a larger z-score represents a better performance.

### Statistical Analysis

Statistical analyses were performed in RStudio [54]. Group differences in demographic, cardiovascular measures, and cognitive measures between the CAD and HC groups were tested using ANCOVAs, controlling for age and sex (and for education in the case of analyses related to cognition). The sex distribution was compared using a Chi-square test. Mean group (CAD vs HC) differences for all quantitative MRI (qMRI) metrics, i.e., *χ*, *R*_2_^∗^, MTsat and *R*_1_ in WS and NWS regions were assessed using ANCOVAs, controlling for age and sex.

To assess whether the association between the cognitive aspects and qMRI is different between CAD and HC groups, a separate linear regression models were used to assess the relationships between each cognitive outcome, including global cognition (MoCA) and the two composite scores (i.e. executive function and verbal episodic memory), and each qMRI metric with group, qMRI-group interaction term, age, sex and education as covariates in WS and NWS regions. Linear regression assumptions were assessed using visual inspection of Q-Q plots and Shapiro–Wilk tests on model residuals. A p-value threshold of 0.05 was used to evaluate residual normality.

To account for multiple comparisons across tests and regions, p-values were adjusted using the Benjamini-Hochberg method (False Discovery Rate, FDR), with a significance threshold set at 0.05.

## Results

Detailed clinical characteristics of the CAD group can be found in Table 1. Briefly, the CAD group consisted of individuals diagnosed with coronary artery disease. Almost half of the CAD group (21) had a history of myocardial infarction, and 65% underwent a percutaneous coronary intervention (i.e., stent), while 24% underwent coronary artery bypass grafting. Individuals with CAD were treated with lipid-lowering medications, beta blockers, aspirin, renin-angiotensin-aldosterone system (RAAS) inhibitors, and calcium channel blockers. Approximately 40% of the HC group had a family history of cardiovascular disease, 15% of them were overweight (BMI > 25), 7% had dyslipidemia and one was treated with a lipid-lowering medication.

**Table 1.** Clinical health profile of the participants.

| Groups | CAD (n = 46) | HC (n = 40) |
| --- | --- | --- |
| **Cardiometabolic risk factors, n (%)** |  |  |
| Type 2 Diabetes | 6 (13) | 0 |
| Hypertension | 23 (50) | 0 |
| Dyslipidemia | 35 (76) | 3 (7) |
| Obesity | 11(24) | 6 (15) |
| **Medication, n (%)** |  |  |
| Beta-Blocker | 21 (46) | 0 |
| Aspirin | 15 (33) | 0 |
| Antilipemic agents (e.g., statins) | 30 (65) | 1 (3) |
| RAAS inhibitor | 12 (26) | 0 |
| Calcium channel blockers | 7 (15) | 0 |
| **Cardiovascular history, n (%)** |  |  |
| Previous MI | 21 (46) | - |
| STEMI | 8 (17) | - |
| NSTEMI | 13 (28) | - |
| Percutaneous coronary intervention (PCI) | 30 (65) | - |
| Coronary artery bypass grafting (CABG) | 11 (24) | - |
| Angina | 13 (28) | - |
| Familial history of cardiovascular disease | 21 (46) | 15 (38) |
|  | **CAD (n = 26)** | - |
| SYNTAX score (mean ± SD) | 13.42 ± 9.45 | - |

MI = Myocardial Infarction; STEMI = ST-elevation Myocardial Infarction; NSTEMI = Non-ST-elevation Myocardial Infarction; RAAS = Renin-Angiotensin-Aldosterone System; SYNTAX = Synergy Between Percutaneous Coronary Intervention with Taxus and Cardiac Surgery; SD = Standard deviation.

Demographic information for both CAD and HC groups can be found in Table 2. There were significant group differences in several parameters, whereby individuals with CAD exhibited higher BMI (p < 0.001; *η*_*p*_^2^ = 0.138), waist circumference (p < 0.001; *η*_*p*_^2^ = 0.18) and body fat mass (p < 0.001; *η*_*p*_^2^ = 0.206) compared to HCs (Table 2). Other demographic parameters such as age, sex, education, systolic and diastolic blood pressure, resting heart rate, and cognition did not differ between groups. The values are reported as mean ± standard deviation in Table 2.

**Table 2.** Demographic data, hemodynamic measures, and cognitive scores in both groups.

| Demographic | CAD (n=46) | HC (n=40) | p-values |
| --- | --- | --- | --- |
| Age (years) | 68.2 ± 8.9 | 65.5 ± 8.0 | 0.145 |
| Sex (M/F) | 38 / 8 | 29 / 11 | 0.386 |
| Education (years) | 16.2 ± 4.5 | 16.0 ± 2.4 | 0.725 |
| BMI (kg/m2) *** | 28.1 ± 4.5 | 26.0 ± 3.9 | < 0.001 |
| Waist Circumference (cm) *** | 102.8 ± 9.4 | 95.1 ± 11.5 | < 0.001 |
| Body Fat (%) *** | 29.0 ± 7.6 | 25.7 ± 7.1 | < 0.001 |
| **Cardiovascular Measures** |  |  |  |
| Systolic BP (mmHg) | 126.8 ± 14.1 | 129.6 ± 14.8 | 0.952 |
| Diastolic BP (mmHg) | 73.5 ± 7.1 | 76.5 ± 9.0 | 0.480 |
| Resting Heart Rate (bpm) | 63.3 ± 10.7 | 68.0 ± 10.4 | 0.146 |
| **Cognition** |  |  |  |
| MoCA | 25.3 ± 2.9 | 26.5 ± 2.1 | 0.092 |
| MMSE | 28.3 ± 1.3 | 28.4 ± 1.4 | 0.692 |
| Executive function (z-score) | -0.19 ± 0.9 | 0.28 ± 0.6 | 0.055 |
| Verbal episodic memory (z-score) | -0.11 ± 1.0 | 0.10 ± 0.9 | 0.370 |

BMI = Body Mass Index; BP = Blood Pressure; MoCA = Montreal Cognitive Assessment; MMSE = Mini-Mental State Examination. The values are in mean ± standard deviation. (Significance Level: \*: $p < 0.05$ , \*\*: $p < 0.01$ , \*\*\*: $p < 0.001$ , \*\*\*\*: $p < 0.0001$ ).

### Lower myelin- and higher iron–sensitive qMRI metrics in CAD

Fig. 3 shows the group differences in total WS region in all qMRI contrasts. In <u>WS</u> areas, the CAD group had significantly higher *χ* in the total WS region (Fig. 3A), as well as in the ACA_MCA and MCA_PCA WS subregions. There were no group differences in *R*_2_^∗^values in any region, though a consistent trend for lower *R*_2_^∗^ in individuals with CAD was observed across the total WS region (Fig. 3B) and all the subregions. The CAD group showed significantly lower *R*_1_ in the total WS region (Figure 3C), and the ACA_MCA and MCA_PCA regions. Furthermore, the CAD group had significantly lower MTsat than HC in total WS (Fig. 3D) and MCA_PCA regions. Supplementary Table 1 includes all group comparisons (including non-significant comparisons) in WS regions for all qMRI contrasts.

**Figure 3.**
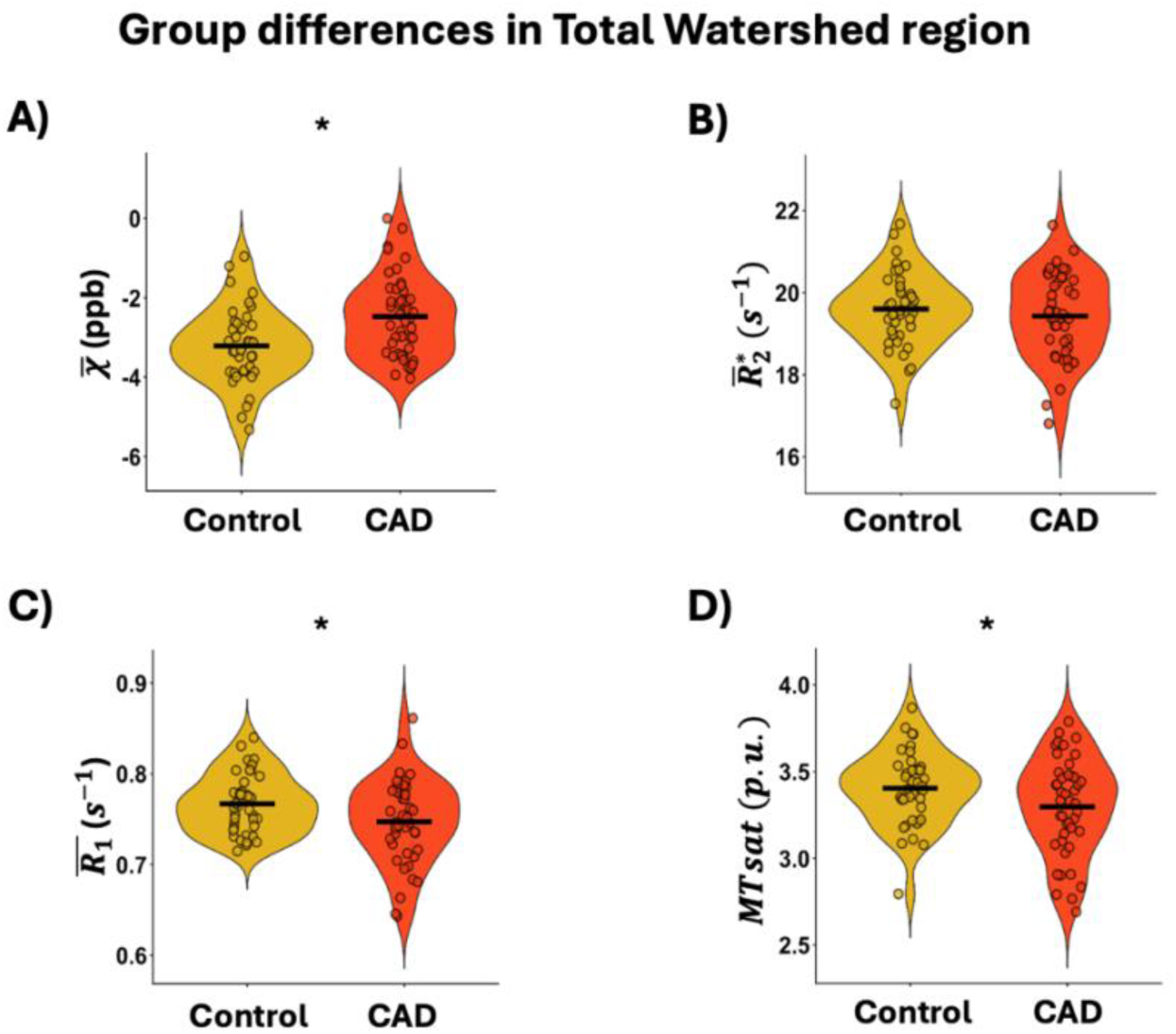
Group comparison of χ, R_2_^∗^, R_1_ and MTsat in the <u>total WS</u> region. **A)** The CAD group showed significantly higher susceptibility than HC. **B)** R_2_^∗^ is nonsignificantly lower in CAD compared to HC. **C)** The CAD group had significantly lower R_1_ than HC. **D)** The CAD group had significantly lower MTsat than HC. The horizontal black line represents the mean value of data points within each violin plot. The unit for χ is part per billion (ppb). The unit for R_2_^∗^ and R_1_ is s^-1^. The unit for MTsat is percent units (p.u.).

In <u>NWS</u> areas, there were no significant group differences for any of the qMRI metrics in any region. It is worth noting however that, similarly to WS regions, all NWS regions showed a trend for higher *χ* and lower *R*_2_^∗^, *R*_1_ and MTsat in individuals with CAD. All statistical results of NWS regions for all contrasts are shown in Supplementary Table 2.

### Cognitive performance is associated with myelin- and iron–sensitive qMRI metrics in CAD

The relationship between qMRI metrics and cognition was assessed, with a focus on whether the CAD and HC groups showed different relationships between WM microstructure and cognitive performance. In <u>WS</u> areas, we observed significant qMRI-group interaction effects between MoCA scores and MTsat, *R*_1_, and *χ*, indicating that the association between MoCA and these qMRI metrics differed between HC and CAD groups (Fig. 4). MTsat showed significant group interactions in the total WS region (pFDR_interaction_ = 0.0192) as well as in the ACA_MCA (pFDR_interaction_ = 0.0439), MCA_PCA (pFDR_interaction_ = 0.0192), and PCA_VB (pFDR_interaction_ = 0.0439) subregions. Post-hoc analyses revealed that, across these regions, Mtsat was not significantly associated with MoCA in HC, whereas higher MTsat was consistently associated with higher MoCA in CAD in the total WS region (p = 0.0001, adjusted-R^2^=0.44), the ACA_MCA (p = 0.0006, adjusted-R^2^=0.38) (Fig. 4A) and the MCA_PCA (p < 0.0001, adjusted-R^2^=0.39) and the PCA_VB (p = 0.0378, adjusted-R^2^=0.22). *R*_1_ showed significant group interactions in the total WS (pFDR_interaction_ = 0.0439) and in the ACA_MCA regions (pFDR_interaction_ = 0.0384). In both regions, *R*_1_ was not significantly associated with MoCA in HC but showed a significant positive association in CAD in the total WS region (p = 0.0021, adjusted-R^2^=0.3) and in the ACA_MCA (p = 0.0090, adjusted-R^2^=0.27) (Fig. 4B). Finally, for *χ*, a significant interaction was observed in the ACA_MCA WS (pFDR_interaction_ = 0.0439). In HC, the relationship between *χ* and MoCA was not significant, whereas in CAD higher *χ* was significantly associated with lower MoCA (p = 0.0314, adjusted-R^2^=0.26) (Fig. 4C). All the group interaction results in WS subregions, including non-significant, are presented in Supplementary Table 3.

**Figure 4.**
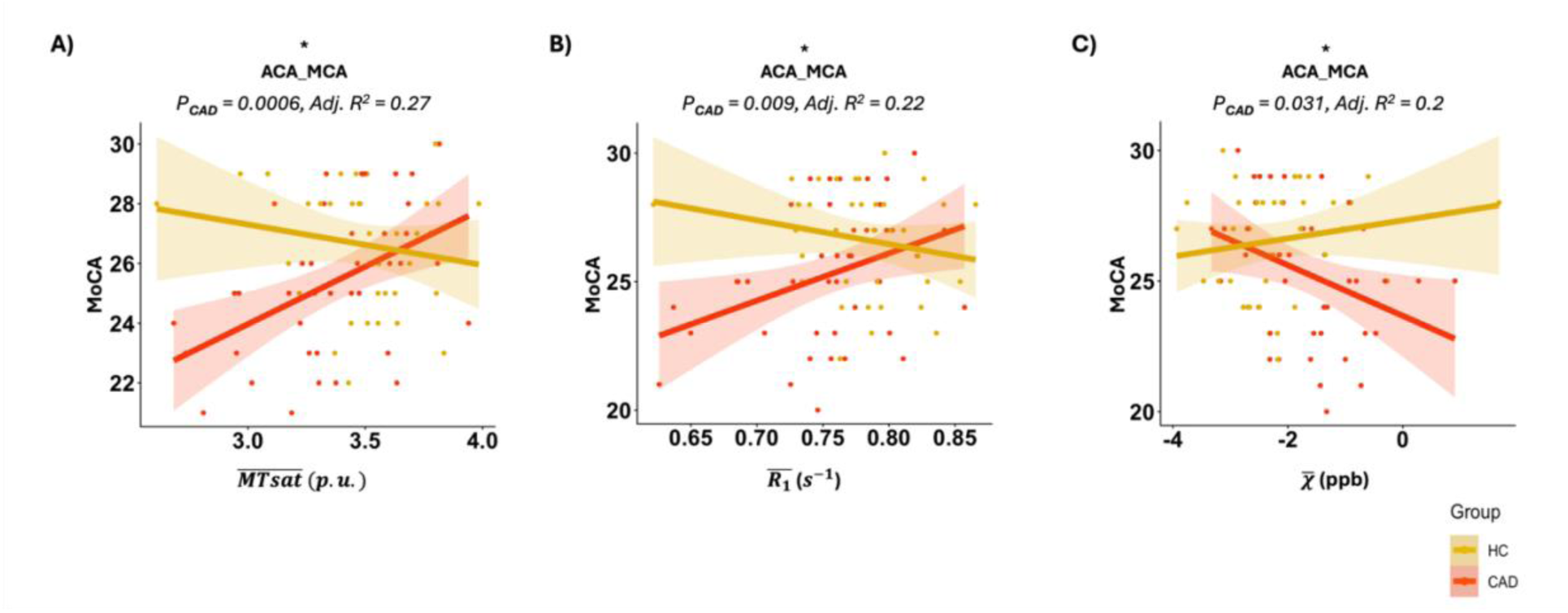
Association between MoCA scores and qMRI metrics in the <u>ACA_MCA</u> WS region in CAD (red-colored line) as a representative example. MoCA score was associated positively with **A)** MTsat, **B)** R_1_and **C)** negatively with χ in the ACA_MCA WS region. The p-value and adjusted-R^2^ values on the figures are corresponding to the relationship in the CAD group. The unit for MTsat is percent units (p.u.). The unit for R_1_ is s^-1^. The unit for χ is part per billion (ppb).

In <u>NWS</u> regions, the interaction models showed several qMRI-group interaction effects (Fig. 5). For MoCA scores, we observed interaction effects for MTsat, *R*_1_ and *R*_2_^∗^. For MTsat, significant interactions were present in the total NWS region (pFDR_interaction_ = 0.0266) and within the ACA (pFDR_interaction_ = 0.0266), the MCA (pFDR_interaction_ < 0.001), and the PCA (pFDR_interaction_ = 0.008) territories. In all regions but the total NWS region, where MTsat was negatively associated with MoCA in HC (p = 0.0495, adjusted-R^2^=0.04), the relationship between MoCA and MTsat was non-significant in HCs. In CAD, there were significant positive relationships between MoCA and MTsat in the ACA (p = 0.0084, adjusted-R^2^=0.3) (Fig. 5A), the MCA (p = 0.0002, adjusted-R^2^=0.42) and the PCA (p = 0.0003, adjusted-R^2^=0.33), though the relationship was marginal, but not significant in the total NWS (p = 0.0698, adjusted-R^2^=0.24). For *R*_1_, significant interactions were observed in ACA (pFDR_interaction_ = 0.0266) (Fig. 5B) and MCA (pFDR_interaction_ = 0.0387). In both territories, R1 was not significantly associated with MoCA in HC, whereas higher R1 was associated with higher MoCA in CAD in the ACA (p = 0.0059, adjusted-R^2^=0.28) and MCA (p = 0.0113, adjusted-R^2^=0.26). For *R*_2_^∗^, a significant interaction was detected in the ACA (pFDR_interaction_ < 0.001), which was specific to the CAD group (p < 0.0001, adjusted-R^2^=0.47) (Fig. 5C).

**Figure 5.**
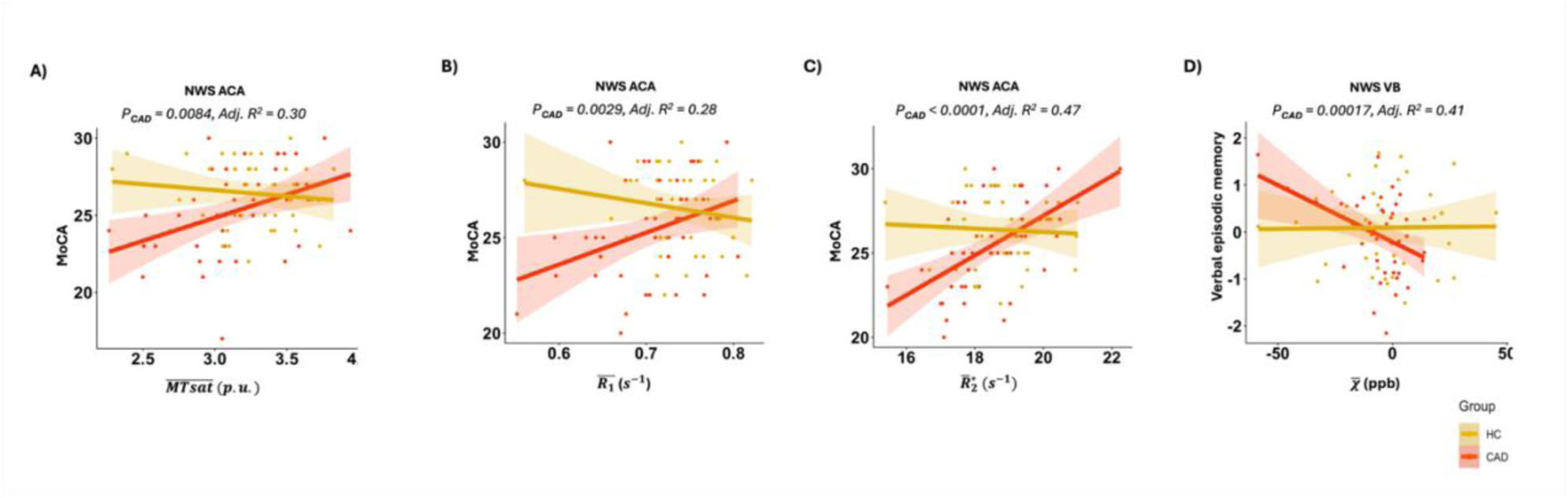
Association between cognition and qMRI metrics in the <u>NWS</u> region in CAD (red-colored line). **A-C)** positive association between MoCA scores and **A)** MTsat, **B)** R_1_ and **C)** R_2_^∗^in NWS ACA territory as a representative example. **D)** X showed negative association with verbal episodic memory in NWS VB territory. The p-value and adjusted-R^2^ values on the figures are corresponding to the relationship in the CAD group. The unit for MTsat is percent units (p.u.). The unit for R_1_ and R_2_^∗^ is s^-1^. The unit for χ is part per billion (ppb). Adj.-R^2^: adjusted-R^2^

For Executive Function, an interaction with *R*_2_^∗^ in VB NWS remained significant after FDR correction (pFDR_interaction_ = 0.0266). In HC, higher *R*_2_^∗^ was significantly associated with lower executive function performance (p = 0.0019, adjusted-R^2^=0.46), while this association was not significant in CAD.

For Verbal Episodic Memory, a significant interaction was observed with *χ* in VB NWS (pFDR_interaction_ = 0.0308). *χ* was not associated with verbal episodic memory in HC, whereas in CAD, higher *χ* was significantly associated with lower verbal episodic memory performance (p = 0.0017, adjusted-R^2^=0.41) (Fig. 5D). The result of all group interaction in NWS subregions, including non-significant, is presented in the Supplementary Table 4.

## Discussion

In this study, we investigated the impact of CAD on NAWM microstructure using advanced quantitative MRI measures sensitive to myelin and iron content. We found that participants with CAD exhibited higher magnetic susceptibility, lower MTsat and lower *R*_1_, specifically within WS regions. This pattern suggests that CAD is associated with both iron accumulation and demyelination in the WS areas, which are known to be especially vulnerable WM regions. Furthermore, this pattern of demyelination and iron deposition was related to cognitive performance uniquely in individuals with CAD: individuals with lower MTsat, lower *R*_1_and higher *χ* showed poorer global cognition (MoCA scores) within WS regions. Similar to WS regions, in NWS regions individuals with lower MTsat and lower *R*_1_ showed poorer global cognition (MoCA scores) and individuals with higher susceptibility showed poorer verbal episodic memory. Together, these findings demonstrate that coronary artery disease contributes to widespread alterations in NAWM microstructure, which are linked to poorer cognition.

### CAD is associated with lower myelin and higher iron in watershed regions

MTsat and R1 results converged, demonstrating reduced myelin across all WS areas in CAD, but not in NWS areas. MTsat is sensitive to macromolecules and since myelin is approximately 80% of macromolecular lipids and 20% proteins [55], it directly reflects the myelin concentration in white matter [56]. Similarly, *R*_1_ is primarily determined by myelin content (∼90%) in white matter, with only a minor contribution from iron (∼10%) [57]. One region, the ACA_MCA, only showed a reduced R1 but not a reduced MTsat in CAD, likely attributable to the fact that while MTsat is more specific to myelin than R1, it may be less sensitive due to signal variability [58]. These results complement previous work demonstrating reduced white matter integrity in CAD [9, 30, 31]. Here we provide additional specificity over previous findings both by relying on myelin-specific qMRI metrics instead of diffusion-based metrics [30, 31], and by identifying the WS areas as the most vulnerable WM areas to microstructural impairments [9].

We also found that participants with CAD exhibited higher magnetic susceptibility in all WS regions relative to healthy controls. In white matter, susceptibility primarily reflects myelin (∼70%) and iron (∼30%) content [57]. Because myelin has negative susceptibility, demyelination leads to less negative (closer to zero) values. Likewise, iron accumulation increases susceptibility. Therefore, our observation of higher susceptibility in CAD is consistent with demyelination alone or combined with iron deposition.

While susceptibility alone cannot distinguish these two mechanisms, the presence of iron deposition can also be inferred from the apparent transverse relaxation rate (*R*_2_^∗^). *R*_2_^∗^ reflects contributions from both myelin (∼52%) and iron (∼48%) but in opposite directions: demyelination lowers *R*_2_^∗^, whereas iron deposition increases it [57]. Because we observed no difference in *R*_2_^∗^ between CAD and control groups, it is likely that both demyelination and iron accumulation are present. If demyelination occurred alone, we would expect to see increased *χ* and lower *R*_2_^∗^. The fact that we did not observe a lower *R*_2_^∗^, suggests that iron accumulation is concomitant with demyelination, increasing the effects of CAD on *χ* and countering the lowering of *R*_2_^∗^ from demyelination with an increased *R*_2_^∗^ from iron.

Therefore, our findings with all three contrasts indicate that, in addition to demyelination, iron deposition may be occurring within the WS regions of individuals with CAD. This pattern could reflect secondary processes such as neuroinflammation [59], microglial activation [60], and increased blood–brain barrier permeability [61], all of which have been associated with cardiovascular disease [62]. However, the trend toward lower *R*_2_^∗^ values in CAD implies that demyelination may be the dominant process, consistent with the observed reductions in MTsat and *R*_1_.

In contrast to the WS regions, NWS white matter did not show significant group differences in any qMRI metric. This is an important observation, as it is consistent with WM microstructural changes being a downstream effect of vascular impairments, since they are in regions defined by their vascular properties, rather than the changes being tract-specific. This finding aligns with previous evidence that WS territories are particularly vulnerable to ischemic injury and hypoperfusion, which are common precursors of both WS infarcts [63] and white matter hyperintensities [64]. Given that CAD is associated with lower cerebral blood flow [22, 65], our findings suggest that compromised perfusion in these regions may underlie the microstructural degradation of NAWM in CAD, though more work is needed however to determine the underlying mechanisms leading to this lower myelin and higher iron content.

### Poorer cognitive performance is associated with lower myelin and higher iron content

The qMRI–group interactions observed here were predominantly driven by associations that were detectable in the CAD group but weak or absent in HC, across both WS and NWS territories. This pattern may reflect restricted variability and/or ceiling effects in the HC group, which showed both less variability in cognitive scores and in qMRI metrics.

These findings align with a putative contribution of vascular pathology to WM injury, in turn leading to poorer cognitive function, which was only detectable in the context of WM injury in CAD [66, 67].

While the cross-sectional nature of this study cannot assess causation, our results are consistent with a negative impact of CAD on WM microstructure, which in turn is associated with poorer cognition. We found lower MTsat and *R*_1_values (reflecting lower myelin content) and higher magnetic susceptibility (reflecting iron deposition) were associated with poorer global cognition (MoCA). Notably, this relationship with demyelination was most evident in the total and the posterior WS regions located at the interface of the MCA and PCA and NWS MCA and ACA territories, highlighting their vulnerability and relevance to cognitive function. These regions are frequently affected by ischemic events, as strokes most often occur in the MCA and then ACA territories [68]. Furthermore, the negative association between MoCA scores and iron deposition which was present in the WS region located between ACA and MCA likely reflects an adverse effect of neuroinflammation and microglial activity on cognition [69].

Executive function showed a significant negative association with *R*_2_^∗^ (Supplementary table 4) in the NWS VB region in the healthy control group. The VB arteries supply crucial regions, including the medial temporal lobe and cerebellum [70]. This finding is consistent with a role for the medial temporal lobe in executive function, such as observed in individuals with Alzheimer’s Disease [71], and with a role of cerebellar WM (i.e. VB arterial territory) integrity in executive function among healthy controls [72]. More work is needed however to understand the mechanistic underpinnings of this association.

Poorer verbal episodic memory was related to higher magnetic susceptibility, i.e. iron deposition, in the NWS VB territory. Woodcock *et. al* [73] demonstrated that in healthy older adults greater microglial activation measured with positron emission tomography was associated with poorer verbal memory in various brain regions including hippocampus, thalamus and temporal lobe which all are supplied by VB arteries [70]. Similarly, Lindgren *et. al* [74] reported that twin pairs with verbal memory deficits exhibited higher uptake of inflammation-sensitive positron emission tomography tracers than those without such deficits in medial temporal lobe, also supplied by VB region. Together, these findings align with our results, suggesting that iron accumulation in CAD could be indicative of neuroinflammatory activity in normal-appearing white matter, which may contribute to poorer verbal episodic memory in these individuals. Studies in CAD using more specific markers of neuroinflammation are however needed to confirm these results.

## Limitations

The primary limitation of this study is limited and sex-unbalanced sample size, which limits our ability to detect more subtle and sex-related effects. Future studies with larger and more sex-balanced cohorts and a wider range of health status and other demographic characteristics are needed to validate our results.

Secondly, future studies should use more precise measures of neuroinflammation such as positron emission tomography [75] techniques alongside with myelin/iron-sensitive qMRI techniques to study the demyelination and neuroinflammation quantitatively for therapeutic purposes.

## Conclusion

This study revealed that coronary artery disease is linked with microstructural alterations in normal-appearing white matter that are most pronounced in watershed regions. These changes reflect primarily demyelination, though accompanied by iron accumulation, a marker of neuroinflammatory processes. The relevance of these WM microstructural alterations is demonstrated by their relationship with cognitive performance across global cognition, as well as executive function and verbal episodic memory. Together, the findings highlight the critical vulnerability of watershed normal-appearing white matter and suggest that maintaining microstructural integrity may help preserve cognitive function in individuals with CAD. Future longitudinal and interventional studies are warranted to determine the mechanisms driving these changes and to identify strategies for neuroprotection in individuals with coronary artery disease.

## Supporting information

Supplementary material is available

## Data availability

Anonymized and defaced data that support the findings of this study will be made openly available on OpenNeuro (https://openneuro.org) within six months of publication. A persistent DOI will be provided upon release.

## Acknowledgments

We would like to thank everyone who contributed to this project: Paule Samson, Thomas Vincent, Julie Lalongé, Hakima Benhalima, Milla Shakleva, Victoria D’Amours, Agathe Godet, Stephanie Beram, Roni Zaks, Robert Hovey, Alexandre Bailey, Catherina Medeiros, Amélie Mainville-Berthiaume and Zineb Rouabah. Thank you also to the laboratories of Dr Louis Bherer and Dr Mathieu Gayda. Lastly, we would like to acknowledge our research participants without whom none of this would have been possible.

## Supplementary material

Supplementary material is available online.

## References

[1] (2025). Cardiovascular diseases (CVDs). https://www.who.int/news-room/fact-sheets/detail/cardiovascular-diseases-(cvds). Accessed 28 Mar 2025.

[2] Mensah GA, Fuster V, Murray CJL, Roth GA, Mensah GA, Abate YH, et al. (2023). Global Burden of Cardiovascular Diseases and Risks, 1990-2022. J Am Coll Cardiol, 82:2350.

[3] Liang X, Huang Y, Han X (2021). Associations between coronary heart disease and risk of cognitive impairment: A meta-analysis. Brain Behav, 11:e02108.

[4] Van Nieuwkerk AC, Delewi R, Wolters FJ, Muller M, Daemen M, Biessels GJ (2023). Cognitive Impairment in Patients with Cardiac Disease: Implications for Clinical Practice. Stroke, 54:2181–2191.

[5] Testai FD, Gorelick PB, Chuang PY, Dai X, Furie KL, Gottesman RF, et al. (2024). Cardiac Contributions to Brain Health: A Scientific Statement from the American Heart Association. Stroke, 55:e425–e438.

[6] Singh-Manoux A, Sabia S, Lajnef M, Ferrie JE, Nabi H, Britton AR, et al. (2008). History of coronary heart disease and cognitive performance in midlife: the Whitehall II study. Eur Heart J, 29:2100–2107.

[7] Madden DJ, Bennett IJ, Song AW (2009). Cerebral white matter integrity and cognitive aging: Contributions from diffusion tensor imaging. Neuropsychol Rev, 19:415–435.

[8] Tuladhar AM, Van Norden AGW, De Laat KF, Zwiers MP, Van Dijk EJ, Norris DG, et al. (2015). White matter integrity in small vessel disease is related to cognition. NeuroImage Clin, 7:518–524.

[9] Tremblay SA, Potvin-Jutras Z, Sabra D, Rezaei A, Sanami S, Gagnon C, et al. (2025). Multivariate white matter microstructure alterations in older adults with coronary artery disease. J Neurosci, e0790252025.

[10] Dadar M, Maranzano J, Ducharme S, Collins DL (2019). White matter in different regions evolves differently during progression to dementia. Neurobiol Aging, 76:71–79.

[11] Huang J, Oh M, Robert C, Huang X, Egle M, Tozer DJ, et al. (2024). Loss of white matter integrity mediates the association between cortical cerebral microinfarcts and cognitive dysfunction: A longitudinal study. J Cereb Blood Flow Metab. doi: 10.1177/0271678X241258563.

[12] Urgesi C, Pavlovic A, Gu Y, Wu D (2022). Association between white matter alterations and domain-specific cognitive impairment in cerebral small vessel disease: A meta-analysis of diffusion tensor imaging. 2022.

[13] Eilaghi A, Kassner A, Sitartchouk I, Francis PL, Jakubovic R, Feinstein A, et al. (2013). Normal-appearing white matter permeability distinguishes poor cognitive performance in processing speed and working memory. Am J Neuroradiol, 34:2119–2124.

[14] Sun Y, Hu W, Hu Y, Qiu Y, Chen Y, Xu Q, et al. (2024). Exploring cognitive related microstructural alterations in normal appearing white matter and deep grey matter for small vessel disease: A quantitative susceptibility mapping study. NeuroImage. doi: 10.1016/j.neuroimage.2024.120790.

[15] Zheng L, Mack WJ, Chui HC, Heflin L, Mungas D, Reed B, et al. (2012). Coronary artery disease is associated with cognitive decline independent of changes on magnetic resonance imaging in cognitively normal elderly adults. J Am Geriatr Soc, 60:499–504.

[16] Khan W, Khlif MS, Mito R, Dhollander T, Brodtmann A (2021). Investigating the microstructural properties of normal-appearing white matter (NAWM) preceding conversion to white matter hyperintensities (WMHs) in stroke survivors. NeuroImage. doi: 10.1016/j.neuroimage.2021.117839.

[17] Sagnier S, Catheline G, Dilharreguy B, Linck PA, Coupé P, Munsch F, et al. (2020). Normal-Appearing White Matter Integrity Is a Predictor of Outcome After Ischemic Stroke. Stroke, 51:449–456.

[18] Kang P, Ying C, Chen Y, Ford AL, An H, Lee JM (2022). Oxygen Metabolic Stress and White Matter Injury in Patients with Cerebral Small Vessel Disease. Stroke, 53:1570–1579.

[19] Torvik A (1984). The Pathogenesis of Watershed Infarcts in the Brain. 1984.

[20] Dogariu OA, Dogariu I, Vasile CM, Berceanu MC, Raicea VC, Albu CV, et al. (2023). Diagnosis and treatment of Watershed strokes: a narrative review. J Med Life, 16:842–850.

[21] Kapasi A, Leurgans SE, James BD, Boyle PA, Arvanitakis Z, Nag S, et al. (2018). Watershed microinfarct pathology and cognition in older persons. Neurobiol Aging, 70:10–17.

[22] Sanami S, Tremblay SA, Rezaei A, Potvin-Jutras Z, Sabra D, Intzandt B, et al. (2025). The Impact of Coronary Artery Disease on Brain Vascular and Metabolic Health: Links to Cognitive Function. Aging Dis, 0.

[23] Möller HE, Bossoni L, Connor JR, Crichton RR, Does MD, Ward RJ, et al. (2019). Iron, Myelin, and the Brain: Neuroimaging Meets Neurobiology. Trends Neurosci, 42:384–401.

[24] Easson K, Gilbert G, Rohlicek CV, Saint-Martin C, Descoteaux M, Deoni SCL, et al. (2022). Altered myelination in youth born with congenital heart disease. Hum Brain Mapp, 43:3545–3558.

[25] Roy B, Vacas S, Ehlert L, Townsley M, Carrier M, Fonarow GC, et al. (2023). Heart Failure-Induced Brain Myelin Changes and Differences Between Sexes. J Neurosci Res, 101:1662–1674.

[26] Liao Z, Dang C, Li M, Bu Y, Han R, Jiang W (2019). Microstructural damage of normal-appearing white matter in subcortical ischemic vascular dementia is associated with Montreal Cognitive Assessment scores. J Int Med Res, 47:5723–5731.

[27] Sagnier S, Catheline G, Dilharreguy B, Linck PA, Coupé P, Munsch F, et al. (2022). Normal-Appearing White Matter Deteriorates over the Year After an Ischemic Stroke and Is Associated with Global Cognition. Transl Stroke Res, 13:716–724.

[28] Deng J, Chen C, Xue S, Su D, Poon WS, Hou H, et al. (2023). Microglia-mediated inflammatory destruction of neuro-cardiovascular dysfunction after stroke. Front Cell Neurosci. doi: 10.3389/fncel.2023.1117218.

[29] Elendu C, Amaechi DC, Elendu TC, Ibhiedu JO, Egbunu EO, Ndam AR, et al. (2023). Stroke and cognitive impairment: understanding the connection and managing symptoms. Ann Med Surg, 85:6057–6066.

[30] Li T, Qin R, Li C, Li L, Wang X, Wang L (2024). Diffusion kurtosis imaging of brain white matter alteration in patients with coronary artery disease based on the TBSS method. Front Aging Neurosci, 16:1301826.

[31] Poirier SE, Suskin NG, Khaw AV, Thiessen JD, Shoemaker JK, Anazodo UC (2024). Probing Evidence of Cerebral White Matter Microstructural Disruptions in Ischemic Heart Disease Before and Following Cardiac Rehabilitation: A Diffusion Tensor MR Imaging Study. J Magn Reson Imaging, 59:2137–2149.

[32] Santiago C, Herrmann N, Swardfager W, Saleem M, Oh PI, Black SE, et al. (2015). White Matter Microstructural Integrity Is Associated with Executive Function and Processing Speed in Older Adults with Coronary Artery Disease. Am J Geriatr Psychiatry, 23:754–763.

[33] Banzett RB, Lansing RW, Evans KC, Shea SA (1996). Stimulus-response characteristics of CO2-induced air hunger in normal subjects. Respir Physiol, 103:19–31.

[34] Xu B, Liu T, Spincemaille P, Prince M, Wang Y (2014). Flow compensated quantitative susceptibility mapping for venous oxygenation imaging. Magn Reson Med, 72:438–445.

[35] Helms G, Dathe H, Kallenberg K, Dechent P (2008). High-resolution maps of magnetization transfer with inherent correction for RF inhomogeneity and T1 relaxation obtained from 3D FLASH MRI. Magn Reson Med, 60:1396–1407.

[36] Chung S, Kim D, Breton E, Axel L (2010). Rapid B1+ mapping using a preconditioning RF pulse with turboFLASH readout. Magn Reson Med, 64:439–446.

[37] Eskildsen SF, Coupé P, Fonov V, Manjón JV, Leung KK, Guizard N, et al. (2012). BEaST: Brain extraction based on nonlocal segmentation technique. NeuroImage, 59:2362–2373.

[38] MINC - Wikibooks, open books for an open world. https://en.wikibooks.org/wiki/MINC. Accessed 23 Oct 2025.

[39] Dadar M, Louis Collins | D (2021). BISON: Brain tissue segmentation pipeline using T 1-weighted magnetic resonance images and a random forest classifier. Magn Reson Med, 85:1881–1894.

[40] Langkammer C, Bredies K, Poser BA, Barth M, Reishofer G, Fan AP, et al. (2015). Fast quantitative susceptibility mapping using 3D EPI and total generalized variation. NeuroImage, 111:622–630.

[41] Dymerska B, Eckstein K, Bachrata B, Siow B, Trattnig S, Shmueli K, et al. (2021). Phase unwrapping with a rapid opensource minimum spanning tree algorithm (ROMEO). Magn Reson Med, 85:2294–2308.

[42] Liu T, Liu J, De Rochefort L, Spincemaille P, Khalidov I, Ledoux JR, et al. (2011). Morphology enabled dipole inversion (MEDI) from a single-angle acquisition: Comparison with COSMOS in human brain imaging. Magn Reson Med, 66:777–783.

[43] Bazin PL, Plessis V, Fan AP, Villringer A, Gauthier CJ (2016). Vessel segmentation from quantitative susceptibility maps for local oxygenation venography. Proc. - Int. Symp. Biomed. Imaging. IEEE Computer Society, 1135–1138.

[44] Straub S, Schneider TM, Emmerich J, Freitag MT, Ziener CH, Schlemmer H-P, et al. (2017). Suitable Reference Tissues for Quantitative Susceptibility Mapping of the Brain. Int Soc Magn Reson Med, 78:204–214.

[45] Alonso-Ortiz E, Levesque IR, Paquin R, Pike GB (2017). Field inhomogeneity correction for gradient echo myelin water fraction imaging. Magn Reson Med, 78:49–57.

[46] Kan H, Kasai H, Arai N, Kunitomo H, Hirose Y, Shibamoto Y (2016). Background field removal technique using regularization enabled sophisticated harmonic artifact reduction for phase data with varying kernel sizes. Magn Reson Imaging, 34:1026–1033.

[47] Pei M, Nguyen TD, Thimmappa ND, Salustri C, Dong F, Cooper MA, et al. (2015). Algorithm for fast monoexponential fitting based on Auto-Regression on Linear Operations (ARLO) of data. Magn Reson Med, 73:843–850.

[48] Tabelow K, Balteau E, Ashburner J, Callaghan MF, Draganski B, Helms G, et al. (2019). hMRI – A toolbox for quantitative MRI in neuroscience and clinical research. NeuroImage, 194:191–210.

[49] Smith SM (2002). Fast robust automated brain extraction. Hum Brain Mapp, 17:143–155.

[50] Liu CF, Hsu J, Xu X, Kim G, Sheppard SM, Meier EL, et al. (2023). Digital 3D Brain MRI Arterial Territories Atlas. Sci Data 2023 101, 10:1–17.

[51] Nasreddine ZS, Phillips NA, Bédirian V, Charbonneau S, Whitehead V, Collin I, et al. (2005). The Montreal Cognitive Assessment, MoCA: a brief screening tool for mild cognitive impairment. J Am Geriatr Soc, 53:695–699.

[52] Delis DC, Kaplan E, Kramer JH (2012). Delis-Kaplan Executive Function System. PsycTESTS Dataset. doi: 10.1037/T15082-000.

[53] Desjardins-Creau L, Berryman N, Minh Vu TT, Villalpando JM, Kergoat MJ, Li KZ, et al. (2014). Physical Functioning Is Associated With Processing Speed and Executive Functions in Community-Dwelling Older Adults. J Gerontol Ser B, 69:837–844.

[54] Posit team (2025). RStudio: Integrated Development Environment for R..

[55] Laule C, Vavasour IM, Kolind SH, Li DKB, Traboulsee TL, Moore GRW, et al. (2007). Magnetic resonance imaging of myelin. Neurotherapeutics, 4:460–484.

[56] Sled JG (2018). Modelling and interpretation of magnetization transfer imaging in the brain. NeuroImage, 182:128–135.

[57] Stüber C, Morawski M, Schäfer A, Labadie C, Wähnert M, Leuze C, et al. (2014). Myelin and iron concentration in the human brain: A quantitative study of MRI contrast. NeuroImage, 93:95–106.

[58] Pietroboni AM, Colombi A, Contarino VE, Russo FML, Conte G, Morabito A, et al. (2023). Quantitative susceptibility mapping of the normal-appearing white matter as a potential new marker of disability progression in multiple sclerosis. Eur Radiol, 33:5368–5377.

[59] Traub J, Frey A, Störk S (2023). Chronic Neuroinflammation and Cognitive Decline in Patients with Cardiac Disease: Evidence, Relevance, and Therapeutic Implications. Life, 13:329.

[60] Möller HE, Bossoni L, Connor JR, Crichton RR, Does MD, Ward RJ, et al. (2019). Iron, Myelin, and the Brain: Neuroimaging Meets Neurobiology. Trends Neurosci, 42:384–401.

[61] Gelosa P, Castiglioni L, Rzemieniec J, Muluhie M, Camera M, Sironi L (2022). Cerebral derailment after myocardial infarct: mechanisms and effects of the signaling from the ischemic heart to brain. J Mol Med, 100:23–41.

[62] Henein MY, Vancheri S, Longo G, Vancheri F (2022). The Role of Inflammation in Cardiovascular Disease. Int J Mol Sci, 23:12906.

[63] El-Gammal TM, Bahnasy WS, Ragab OAA, AL-Malt AM (2018). Cerebral border zone infarction: an etiological study. Egypt J Neurol Psychiatry Neurosurg, 54:1–6.

[64] Wang Y, Liu G, Hong D, Chen F, Ji X, Cao G (2016). White matter injury in ischemic stroke. Prog Neurobiol, 141:45–60.

[65] Anazodo UC, Shoemaker JK, Suskin N, Ssali T, Wang DJJ, St. Lawrence KS (2016). Impaired cerebrovascular function in coronary artery disease patients and recovery following cardiac rehabilitation. Front Aging Neurosci. doi: 10.3389/fnagi.2015.00224.

[66] Moroni F, Ammirati E, Rocca MA, Filippi M, Magnoni M, Camici PG (2018). Cardiovascular disease and brain health: Focus on white matter hyperintensities. Int J Cardiol Heart Vasc, 19:63–69.

[67] Rundek T, Tolea M, Ariko T, Fagerli EA, Camargo CJ (2022). Vascular Cognitive Impairment (VCI). Neurotherapeutics, 19:68–88.

[68] Nichols L, Bridgewater JC, Wagner NB, Karivelil M, Koelmeyer H, Goings D, et al. (2021). Where in the Brain do Strokes Occur? A Pilot Study and Call for Data. Clin Med Res, 19:110–115.

[69] Ownby RL (2010). Neuroinflammation and Cognitive Aging. Curr Psychiatry Rep, 12:39–45.

[70] Benjamin R, Lui F (2025). Vertebrobasilar Insufficiency. StatPearls.

[71] Oosterman JM, Oosterveld S, Rikkert MGO, Claassen JA, Kessels RPC (2012). Medial temporal lobe atrophy relates to executive dysfunction in Alzheimer’s disease. Int Psychogeriatr, 24:1474–1482.

[72] Urbini N, McNabb CB, Jones DK, Hedge C, Messaritaki E, Laguna PL, et al. (2025). The cognitive cerebellum: linking microstructure to cognitive functions in a healthy population. NeuroImage, 317:121356.

[73] Woodcock EA, Hillmer AT, Sandiego CM, Maruff P, Carson RE, Cosgrove KP, et al. (2021). Acute neuroimmune stimulation impairs verbal memory in adults: A PET brain imaging study. Brain Behav Immun, 91:784–787.

[74] Lindgren N, Tuisku J, Vuoksimaa E, Helin S, Karrasch M, Marjamäki P, et al. (2020). Association of neuroinflammation with episodic memory: a [11C]PBR28 PET study in cognitively discordant twin pairs. Brain Commun. doi: 10.1093/braincomms/fcaa024.

[75] Kreisl WC, Kim M-J, Coughlin JM, Henter ID, Owen DR, Innis RB (2020). PET Imaging of Neuroinflammation in Neurological Disorders. Lancet Neurol, 19:940–950.

