## Supplementary material is available for "Coronary artery disease is linked with demyelination and iron deposition in white matter watershed areas"

**Supplementary Table 1 - Group comparison of qMRI metrics in watershed regions. (Significance level in the interaction term: *: p < 0.05, **: p < 0.01, ***: p < 0.001)**

| **Group comparison** | | | | |
| --- | --- | --- | --- | --- |
|  | **CAD (n=46)** | **HC (n=40)** | **P_FDR_** | **p.**$\boldsymbol{\eta}^{\mathbf{2}}$ |
| **X (ppm)** | | | | |
| **Total ws *** | **-0.0025 ± 0.001** | **-0.0032 ± 0.001** | **0.016** | **0.14** |
| **ACA_MCA *** | **-0.0017 ± 0.001** | **-0.0023 ± 0.001** | **0.016** | **0.11** |
| **MCA_PCA *** | **-0.0034 ± 0.001** | **-0.0042 ± 0.001** | **0.019** | **0.09** |
| **PCA_VB** | **-0.0038 ± 0.003** | **0.0054 ± 0.004** | **0.06** | **0.05** |
| **R_2_* (Hz)** | | | | |
| **Total ws** | **19.43 ± 1.046** | **19.61 ± 0.911** | **0.442** | **0.01** |
| **ACA_MCA** | **18.85 ± 1.18** | **19.02 ± 0.99** | **0.452** | **0.01** |
| **MCA_PCA** | **20.11 ± 1.09** | **20.28 ± 0.95** | **0.451** | **0.01** |
| **PCA_VB** | **19.73 ± 1.22** | **20 ± 1.07** | **0.338** | **0.01** |
| **R_1_ (Hz)** | | | | |
| **Total ws *** | **0.747 ± 0.046** | **0.767 ± 0.033** | **0.032** | **0.08** |
| **ACA_MCA *** | **0.761 ± 0.046** | **0.783 ± 0.036** | **0.019** | **0.09** |
| **MCA_PCA *** | **0.729 ± 0.043** | **0.751 ± 0.031** | **0.016** | **0.11** |
| **PCA_VB** | **0.687 ± 0.079** | **0.713 ± 0.058** | **0.138** | **0.04** |
| **MTsat (p.u.)** | | | | |
| **Total ws *** | **3.29 ± 0.28** | **3.41 ± 0.22** | **0.034** | **0.07** |
| **ACA_MCA** | **3.41 ± 0.28** | **3.48 ± 0.24** | **0.167** | **0.03** |
| **MCA_PCA *** | **3.24 ± 0.26** | **3.36 ± 0.21** | **0.019** | **0.09** |
| **PCA_VB** | **2.64 ± 0.42** | **2.74 ± 0.41** | **0.297** | **0.02** |

***P_FDR_: FDR-corrected p-value.***

***p.***$\boldsymbol{\eta}^{\boldsymbol{2}}$***: partial*** $\eta^{2}$

**Supplementary Table 2 - Group comparison of qMRI metrics in non-watershed regions. (Significance level in the interaction term: *: p < 0.05, **: p < 0.01, ***: p < 0.001)**

| **Group comparison** | | | | |
| --- | --- | --- | --- | --- |
|  | **CAD (n=46)** | **HC (n=40)** | **P_FDR_** | **p.**$\boldsymbol{\eta}^{\mathbf{2}}$ |
| **X (ppm)** | | | | |
| **Total non-ws** | **-0.0035 ± 0.0042** | **-0.0036 ± 0.0028** | **0.12** | **0.05** |
| **non-ws ACA** | **-0.0032 ± 0.0015** | **-0.0041 ± 0.0016** | **0.086** | **0.08** |
| **non-ws MCA** | **-0.0012 ± 0.0011** | **-0.0016 ± 0.0014** | **0.20** | **0.02** |
| **non-ws PCA** | **-0.0044 ± 0.002** | **-0.0054 ± 0.002** | **0.09** | **0.07** |
| **Non-ws VB** | **-0.005 ± 0.012** | **-0.0033 ± 0.016** | **0.6** | **0.0001** |
| **R_2_* (Hz)** | | | | |
| **Total non-ws** | **18.48 ± 2.4** | **18.3 ± 1.98** | **0.10** | **0.06** |
| **non-ws ACA** | **18.55 ± 1.3** | **18.79 ± 1.19** | **0.39** | **0.01** |
| **non-ws MCA** | **19.44 ± 1.16** | **19.75 ± 1.01** | **0.21** | **0.02** |
| **non-ws PCA** | **19.75 ± 1.03** | **20.14 ± 0.99** | **0.13** | **0.04** |
| **Non-ws VB** | **16.16 ± 4.67** | **14.53 ± 4.74** | **0.17** | **0.03** |
| **R_1_ (Hz)** | | | | |
| **Total non-ws** | **0.14 ± 0.02** | **0.15 ± 0.017** | **0.17** | **0.03** |
| **non-ws ACA *** | **0.71 ± 0.05** | **0.74 ± 0.04** | **0.04** | **0.12** |
| **non-ws MCA** | **0.7 ± 0.06** | **0.73 ± 0.04** | **0.065** | **0.08** |
| **non-ws PCA** | **0.69 ± 0.064** | **0.71 ± 0.051** | **0.09** | **0.05** |
| **Non-ws VB** | **0.42 ± 0.38** | **0.39 ± 0.5** | **0.75** | **0.0002** |
| **MTsat (p.u.)** | | | | |
| **Total non-ws** | **3.35 ± 0.49** | **3.5 ± 0.41** | **0.17** | **0.04** |
| **non-ws ACA** | **3.22 ± 0.35** | **3.28 ± 0.26** | **0.37** | **0.01** |
| **non-ws MCA** | **3.13 ± 0.33** | **3.25 ± 0.26** | **0.1** | **0.06** |
| **non-ws PCA** | **3.1 ± 0.36** | **3.2 ± 0.31** | **0.17** | **0.03** |
| **non-ws VB** | **3.94 ± 0.91** | **4.28 ± 0.82** | **0.17** | **0.04** |

***P_FDR_: FDR-corrected p-value.***

***p.***$\boldsymbol{\eta}^{\boldsymbol{2}}$***: partial*** $\eta^{2}$***.***

**Supplementary Table 3 - Association of cognition and qMRI metrics in watershed regions. (Significance level in the interaction term: *: p < 0.05, **: p < 0.01, ***: p < 0.001)**

| **Relationship with cognitive domains** | | | | | |
| --- | --- | --- | --- | --- | --- |
|  | **pFDR_interaction_** | **p_CAD_** | **Adj.** $\mathbf{R}^{\mathbf{2}}$**_CAD_** | **p_HC_** | **Adj.** $\mathbf{R}^{\mathbf{2}}$**_HC_** |
| **Verbal episodic memory** | | | | | |
| **X (ppm)** |  | | | | |
| **Total ws** | 0.921 | 0.032 | 0.304 | 0.038 | 0.358 |
| **ACA_MCA** | 0.921 | 0.132 | 0.245 | 0.089 | 0.327 |
| **MCA_PCA** | 0.695 | 0.109 | 0.302 | 0.047 | 0.366 |
| **PCA_VB** | 0.979 | 0.836 | 0.181 | 0.818 | 0.323 |
| **R_2_* (Hz)** |  | | | | |
| **Total ws** | 0.921 | 0.481 | 0.196 | 0.021 | 0.377 |
| **ACA_MCA** | 0.921 | 0.625 | 0.191 | 0.019 | 0.382 |
| **MCA_PCA** | 0.921 | 0.921 | 0.546 | 0.196 | 0.443 |
| **PCA_VB** | 0.921 | 0.921 | 0.208 | 0.203 | 0.454 |
| **R_1_ (Hz)** |  | | | | |
| **Total ws** | 0.921 | 0.09 | 0.267 | 0.135 | 0.355 |
| **ACA_MCA** | 0.921 | 0.03 | 0.297 | 0.077 | 0.375 |
| **MCA_PCA** | 0.647 | 0.542 | 0.27 | 0.168 | 0.354 |
| **PCA_VB** | 0.921 | 0.478 | 0.278 | 0.758 | 0.361 |
| **MTsat (p.u.)** |  | | | | |
| **Total ws** | 0.912 | 0.087 | 0.274 | 0.064 | 0.395 |
| **ACA_MCA** | 0.921 | 0.064 | 0.275 | 0.087 | 0.351 |
| **MCA_PCA** | 0.912 | 0.175 | 0.224 | 0.082 | 0.391 |
| **PCA_VB** | 0.489 | 0.038 | 0.319 | 0.927 | 0.293 |
| **Executive function** | | | | | |
| **X (ppm)** |  |  |  | | |
| **Total ws** | 0.77 | 0.191 | 0.596 | 0.55 | 0.489 |
| **ACA_MCA** | 0.692 | 0.562 | 0.433 | 0.945 | 0.353 |
| **MCA_PCA** | 0.565 | 0.149 | 0.545 | 0.757 | 0.412 |
| **PCA_VB** | 0.921 | < 0.001 | 0.612 | 0.474 | 0.314 |
| **R_2_* (Hz)** |  | | | | |
| **Total ws** | 0.489 | 0.562 | 0.433 | 0.079 | 0.518 |
| **ACA_MCA** | 0.489 | 0.149 | 0.545 | 0.161 | 0.491 |
| **MCA_PCA** | 0.565 | < 0.001 | 0.612 | 0.036 | 0.546 |
| **PCA_VB** | 0.135 | 0.262 | 0.572 | 0.029 | 0.554 |
| **R_1_ (Hz)** |  | | | | |
| **Total ws** | 0.565 | < 0.001 | 0.612 | 0.028 | 0.476 |
| **ACA_MCA** | 0.565 | 0.01 | 0.602 | 0.23 | 0.447 |
| **MCA_PCA** | 0.645 | < 0.001 | 0.62 | 0.006 | 0.517 |
| **PCA_VB** | 0.912 | 0.1 | 0.497 | 0.177 | 0.464 |
| **MTsat (p.u.)** |  | | | | |
| **Total ws** | 0.599 | 0.149 | 0.545 | 0.91 | 0.432 |
| **ACA_MCA** | 0.565 | 0.122 | 0.557 | 0.965 | 0.432 |
| **MCA_PCA** | 0.912 | 0.037 | 0.484 | 0.189 | 0.353 |
| **PCA_VB** | 0.599 | 0.502 | 0.462 | 0.446 | 0.335 |
| **MoCA** | | | | | |
| **X (ppm)** |  |  |  | | |
| **Total ws** | 0.119 | 0.008 | 0.35 | 0.761 | -0.011 |
| **ACA_MCA *** | 0.044 | 0.031 | 0.255 | 0.085 | 0.061 |
| **MCA_PCA** | 0.565 | 0.004 | 0.364 | 0.471 | 0.005 |
| **PCA_VB** | 0.692 | 0.106 | 0.159 | 0.68 | 0.008 |
| **R_2_* (Hz)** |  | | | | |
| **Total ws** | 0.599 | 0.039 | 0.22 | 0.652 | -0.009 |
| **ACA_MCA** | 0.565 | 0.042 | 0.217 | 0.788 | -0.01 |
| **MCA_PCA** | 0.912 | 0.275 | 0.176 | 0.683 | -0.011 |
| **PCA_VB** | 0.135 | < 0.001 | 0.35 | 0.768 | -0.014 |
| **R_1_ (Hz)** |  | | | | |
| **Total ws *** | 0.044 | 0.002 | 0.301 | 0.465 | -0.025 |
| **ACA_MCA *** | 0.038 | 0.009 | 0.268 | 0.174 | 0.025 |
| **MCA_PCA** | 0.224 | 0.001 | 0.328 | 0.884 | -0.033 |
| **PCA_VB** | 0.489 | 0.057 | 0.266 | 0.702 | -0.021 |
| **MTsat (p.u.)** |  |  |  | | |
| **Total ws** | 0.018 | < 0.001 | 0.436 | 0.341 | 0.064 |
| **ACA_MCA *** | 0.044 | 0.001 | 0.38 | 0.739 | 0.002 |
| **MCA_PCA *** | 0.019 | < 0.001 | 0.39 | 0.506 | 0.048 |
| **PCA_VB *** | 0.044 | 0.038 | 0.221 | 0.065 | 0.064 |

***pFDR_interaction_: FDR-corrected p-value of the interaction between Group and qMRI metrics.***

***p_CAD_ & Adj.*** $\boldsymbol{R}^{\boldsymbol{2}}$***_CAD_: p-value and adjusted*** $R^{2}$ ***of the cognition-qMRI linear relationship for the CAD group.***

***p_HC_ & Adj.*** $\boldsymbol{R}^{\boldsymbol{2}}$***_HC_: p-value and adjusted*** $R^{2}$ ***of the cognition-qMRI linear relationship for the HC group.***

**Supplementary Table 4 - Association between cognition and qMRI metrics in non-watershed regions. (Significance level in the interaction term: *: p < 0.05, **: p < 0.01, ***: p < 0.001)**

| **Relationship with cognitive domains** | | | | | | |
| --- | --- | --- | --- | --- | --- | --- |
|  | **pFDR_interaction_** | | **p_CAD_** | **Adj.** $\mathbf{R}^{\mathbf{2}}$**_CAD_** | **p_HC_** | **Adj.** $\mathbf{R}^{\mathbf{2}}$**_HC_** |
| **Verbal episodic memory** | | | | | | |
| **X (ppm)** |  | | | | | |
| **Total non-ws** | 0.798 | 0.592 | | 0.25 | 0.239 | 0.327 |
| **non-ws ACA** | 0.904 | 0.306 | | 0.197 | 0.942 | 0.315 |
| **non-ws MCA** | 0.109 | 0.202 | | 0.239 | 0.096 | 0.353 |
| **non-ws PCA** | 0.25 | 0.199 | | 0.25 | 0.296 | 0.326 |
| **Non-ws VB** | 0.735 | 0.481 | | 0.158 | 0.046 | 0.368 |
| **R_2_* (Hz)** |  | | | | | |
| **Total non-ws** | 0.904 | 0.306 | | 0.197 | 0.942 | 0.315 |
| **non-ws ACA** | 0.109 | 0.202 | | 0.239 | 0.096 | 0.353 |
| **non-ws MCA** | 0.25 | 0.199 | | 0.25 | 0.296 | 0.326 |
| **non-ws PCA** | 0.735 | 0.481 | | 0.158 | 0.046 | 0.368 |
| **Non-ws VB** | 0.965 | 0.435 | | 0.2 | 0.019 | 0.38 |
| **R_1_ (Hz)** |  | | | | | |
| **Total non-ws** | 0.25 | 0.199 | | 0.25 | 0.296 | 0.326 |
| **non-ws ACA** | 0.735 | 0.481 | | 0.158 | 0.046 | 0.368 |
| **non-ws MCA** | 0.965 | 0.435 | | 0.2 | 0.019 | 0.38 |
| **non-ws PCA** | 0.965 | 0.158 | | 0.259 | 0.2 | 0.314 |
| **Non-ws VB** | 0.965 | 0.039 | | 0.296 | 0.071 | 0.388 |
| **MTsat (p.u.)** |  | | | | | |
| **Total non-ws** | 0.109 | 0.202 | | 0.239 | 0.096 | 0.353 |
| **non-ws ACA** | 0.25 | 0.199 | | 0.25 | 0.296 | 0.326 |
| **non-ws MCA** | 0.735 | 0.481 | | 0.158 | 0.046 | 0.368 |
| **non-ws PCA** | 0.965 | 0.435 | | 0.2 | 0.019 | 0.38 |
| **Non-ws VB** | 0.965 | 0.158 | | 0.259 | 0.2 | 0.314 |
| **Executive function** | | | | | | |
| **X (ppm)** |  | | | | | |
| **Total non-ws** | 0.478 | 0.228 | | 0.502 | 0.564 | 0.434 |
| **non-ws ACA** | 0.965 | 0.733 | | 0.416 | 0.439 | 0.34 |
| **non-ws MCA** | 0.735 | 0.127 | | 0.599 | 0.396 | 0.439 |
| **non-ws PCA** | 0.586 | 0.022 | | 0.526 | 0.262 | 0.326 |
| **Non-ws VB** | 0.374 | 0.057 | | 0.626 | 0.99 | 0.353 |
| **R_2_* (Hz)** |  | | | | | |
| **Total non-ws** | 0.965 | 0.733 | | 0.416 | 0.439 | 0.34 |
| **non-ws ACA** | 0.735 | 0.127 | | 0.599 | 0.396 | 0.439 |
| **non-ws MCA** | 0.586 | 0.022 | | 0.526 | 0.262 | 0.326 |
| **non-ws PCA** | 0.374 | 0.057 | | 0.626 | 0.99 | 0.353 |
| **Non-ws VB** | 0.068 | 0.073 | | 0.474 | 0.076 | 0.512 |
| **R_1_ (Hz)** |  | | | | | |
| **Total non-ws** | 0.586 | 0.022 | | 0.526 | 0.262 | 0.326 |
| **non-ws ACA** | 0.374 | 0.057 | | 0.626 | 0.99 | 0.353 |
| **non-ws MCA** | 0.068 | 0.073 | | 0.474 | 0.076 | 0.512 |
| **non-ws PCA** | 0.252 | 0.098 | | 0.565 | 0.536 | 0.438 |
| **Non-ws VB** | 0.739 | 0.009 | | 0.528 | 0.147 | 0.45 |
| **MTsat (p.u.)** |  | | | | | |
| **Total non-ws** | 0.735 | 0.127 | | 0.599 | 0.396 | 0.439 |
| **non-ws ACA** | 0.586 | 0.022 | | 0.526 | 0.262 | 0.326 |
| **non-ws MCA** | 0.374 | 0.057 | | 0.626 | 0.99 | 0.353 |
| **non-ws PCA** | 0.068 | 0.073 | | 0.474 | 0.076 | 0.512 |
| **Non-ws VB** | 0.252 | 0.098 | | 0.565 | 0.536 | 0.438 |
| **MoCA** | | | | | | |
| **X (ppm)** |  | | | | | |
| **Total non-ws** | 0.904 | 0.398 | | 0.241 | 0.288 | -0.009 |
| **non-ws ACA** | 0.367 | 0.208 | | 0.188 | 0.387 | -0.001 |
| **non-ws MCA *** | 0.027 | 0.07 | | 0.244 | 0.05 | 0.041 |
| **non-ws PCA** | 0.25 | 0.485 | | 0.215 | 0.095 | 0.035 |
| **Non-ws VB** | 0.965 | 0.649 | | 0.131 | 0.773 | -0.006 |
| **R_2_* (Hz)** |  | | | | | |
| **Total non-ws** | 0.367 | 0.208 | | 0.188 | 0.387 | -0.001 |
| **non-ws ACA *** | 0.027 | 0.07 | | 0.244 | 0.05 | 0.041 |
| **non-ws MCA** | 0.25 | 0.485 | | 0.215 | 0.095 | 0.035 |
| **non-ws PCA** | 0.965 | 0.649 | | 0.131 | 0.773 | -0.006 |
| **Non-ws VB **** | 0.001 | < 0.001 | | 0.474 | 0.197 | 0.009 |
| **R_1_ (Hz)** |  | | | | | |
| **Total non-ws** | 0.25 | 0.485 | | 0.215 | 0.095 | 0.035 |
| **non-ws ACA** | 0.965 | 0.649 | | 0.131 | 0.773 | -0.006 |
| **non-ws MCA **** | 0.001 | < 0.001 | | 0.474 | 0.197 | 0.009 |
| **non-ws PCA *** | 0.027 | 0.008 | | 0.293 | 0.189 | 0.04 |
| **Non-ws VB *** | 0.027 | 0.006 | | 0.284 | 0.15 | 0.017 |
| **MTsat (p.u.)** |  | | | | | |
| **Total non-ws *** | 0.027 | 0.07 | | 0.244 | 0.05 | 0.041 |
| **non-ws ACA** | 0.25 | 0.485 | | 0.215 | 0.095 | 0.035 |
| **non-ws MCA** | 0.965 | 0.649 | | 0.131 | 0.773 | -0.006 |
| **non-ws PCA **** | 0.001 | < 0.001 | | 0.474 | 0.197 | 0.009 |
| **Non-ws VB *** | 0.027 | 0.008 | | 0.293 | 0.189 | 0.04 |

***pFDR_interaction_: FDR-corrected p-value of the interaction between Group and qMRI metrics.***

***p_CAD_ & Adj.*** $\boldsymbol{R}^{\boldsymbol{2}}$***_CAD_: p-value and adjusted*** $R^{2}$ ***of the cognition-qMRI linear relationship for the CAD group.***

***p_HC_ & Adj.*** $\boldsymbol{R}^{\boldsymbol{2}}$***_HC_: p-value and adjusted*** $R^{2}$ ***of the cognition-qMRI linear relationship for the HC group.***
